# Evolutionary dynamics of recent selection for enhanced social cognition

**DOI:** 10.1101/425215

**Authors:** Sara E. Miller, Andrew W. Legan, Michael Henshaw, Katherine L. Ostevik, Kieran Samuk, Floria M. K. Uy, H. Kern Reeve, Michael J. Sheehan

## Abstract

Cognitive abilities can vary dramatically among species though little is known about the dynamics of cognitive evolution. Here we demonstrate that recent evolution of visual individual recognition in the paper wasp *Polistes fuscatus* is the target of arguably the strongest positive selective pressure in the species’ recent history. The most extreme selective sweeps in *P. fuscatus* are associated with genes known to be involved in long-term memory formation, mushroom body development and visual processing – all traits that have recently evolved in association with individual recognition. Cognitive evolution appears to have been driven initially by selection on standing variation in perceptual traits followed by both hard and soft sweeps on learning and memory. Evolutionary modeling reveals that intense selection as observed in *P. fuscatus* is likely the norm during the early stages of cognitive evolution. These data provide insight into the dynamics of cognition evolution demonstrating that social selection for increased intelligence can lead to rapid multi-genic adaptation of enhanced recognition abilities.

## Main Text

Cognition is arguably among the most complex animal traits and has been instrumental in the ecological and evolutionary success of disparate lineages (*1*). Despite intense interest, the evolutionary dynamics that give rise to novel and elaborated cognitive abilities is poorly understood. As with complex morphological traits, intra-specific variation in cognitive abilities is a heritable quantitative trait (*2*, *3*). Furthermore, genome wide association studies in humans have identified hundreds of loci associated with traits such as educational attainment (*4*), suggesting that evolution of enhanced cognition may be a highly polygenic and gradual process. Comparative genomic analyses have identified signatures of positive selection on cognition-associated genes (*5*–*7*), though the deep time-scale of such analyses obscures the dynamics of cognitive evolution. Population genomic scans have detected evidence of recent selection on the nervous system and cognition-associated genes in species such as humans and great-tits (*8*, *9*), though the links to specific cognitive abilities have not been clear.

The recent evolution of visual individual recognition in the northern paper wasp (*10*), *Polistes fuscatus*, provides an unusual opportunity to examine the mode and tempo of cognitive evolution (Fig 1). In *P. fuscatus*, nests are initiated each spring by one or more mated females (*11*). Individual recognition mediates dominance interactions among nest foundresses (*10*) and has been associated with complex social cognition in *P. fuscatus* including highly robust social memories (*12*), tracking others’ contribution to work and egg laying (*13*), and specialized processing of conspecific facial images (*14*). There is no evidence of major ecological differentiation between *P. fuscatus* and closely related paper wasps, which co-occur throughout much of their range often nesting on the same buildings and capturing the same prey species (*11*). The clearest difference between *P. fuscatus* and its relatives is the elaboration of social cognition.

**Figure 1.**
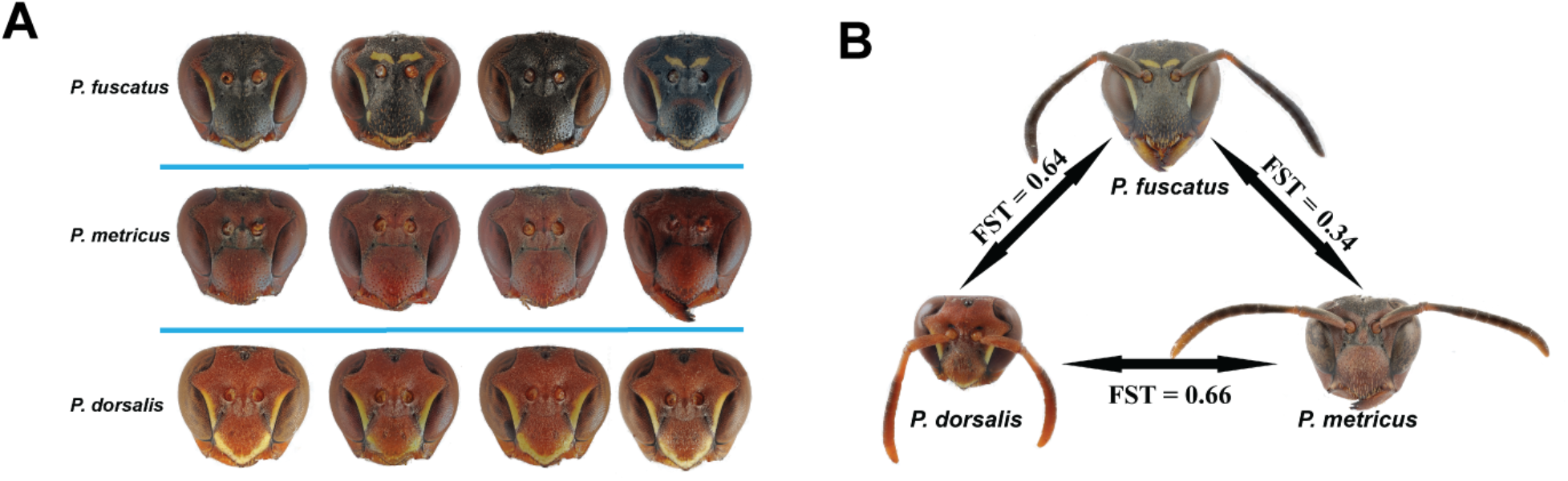
(A) The range of variation in facial coloration within female wasps from a single nest for *P. fuscatus*, *P. metricus*, and *P. dorsalis*. *P. fuscatus* displays greater phenotypic variation in facial coloration than the other species. Antennae have been removed. (B) Pairwise F_ST_ among *P. fuscatus*, *P. metricus*, and *P. dorsalis* show limited differentiation between species. F_ST_ was calculated with all sequences aligned to the *P. fuscatus* genome.

To address the mode and tempo of cognitive evolution, we assembled and annotated high-quality *de novo* genomes for *P. fuscatus* and two close relatives which lack individual recognition, *P. metricus* and *P. dorsalis* (Fig. 1). The three genomes have similar compositions and are comparable to other previously sequenced *Polistes* wasp genomes (Fig S1-S3, Table S1-2, *20*, *21*). We re-sequenced 40 *P. fuscatus*, 19 *P. metricus*, and 18 *P. dorsalis* wasps at an average coverage of 10.6x (Fig. S4-5, Table S3). Pairwise F_ST_ calculations between *P. fuscatus*, *P. metricus*, and *P. dorsalis* shows only moderate genetic differentiation among species (Fig. 1B, Fig. S4), supporting the recent divergence of *P. fuscatus*. Linkage disequilibrium (LD) decays rapidly in all three species (Fig. S5), consistent with the unusually high recombination rates reported in other social hymenopterans (*17*). Recent divergence and rapid LD decay in *P. fuscatus* make it possible to detect selection with exceptionally high precision.

We conducted a genome-wide scan for selective sweeps in *P. fuscatus* identifying 138 peaks of CLR (composite likelihood ratio) consistent with recent selective sweeps (Fig. 2A). Sweeps were narrow (Median 2,843 bp; Range 100-56,811 bp), associated with 183 genes (median = 1 gene/sweep, range: 0-8 genes) and distributed throughout the genome. We were able to further pinpoint candidate regions under selection by calculating sequence differentiation from a reconstructed ancestral genome and plotting this in the same windows (Fig. 2B). These findings indicate that paper wasps provide an unusually tractable system for identifying targets of recent natural selection.

**Figure 2.**
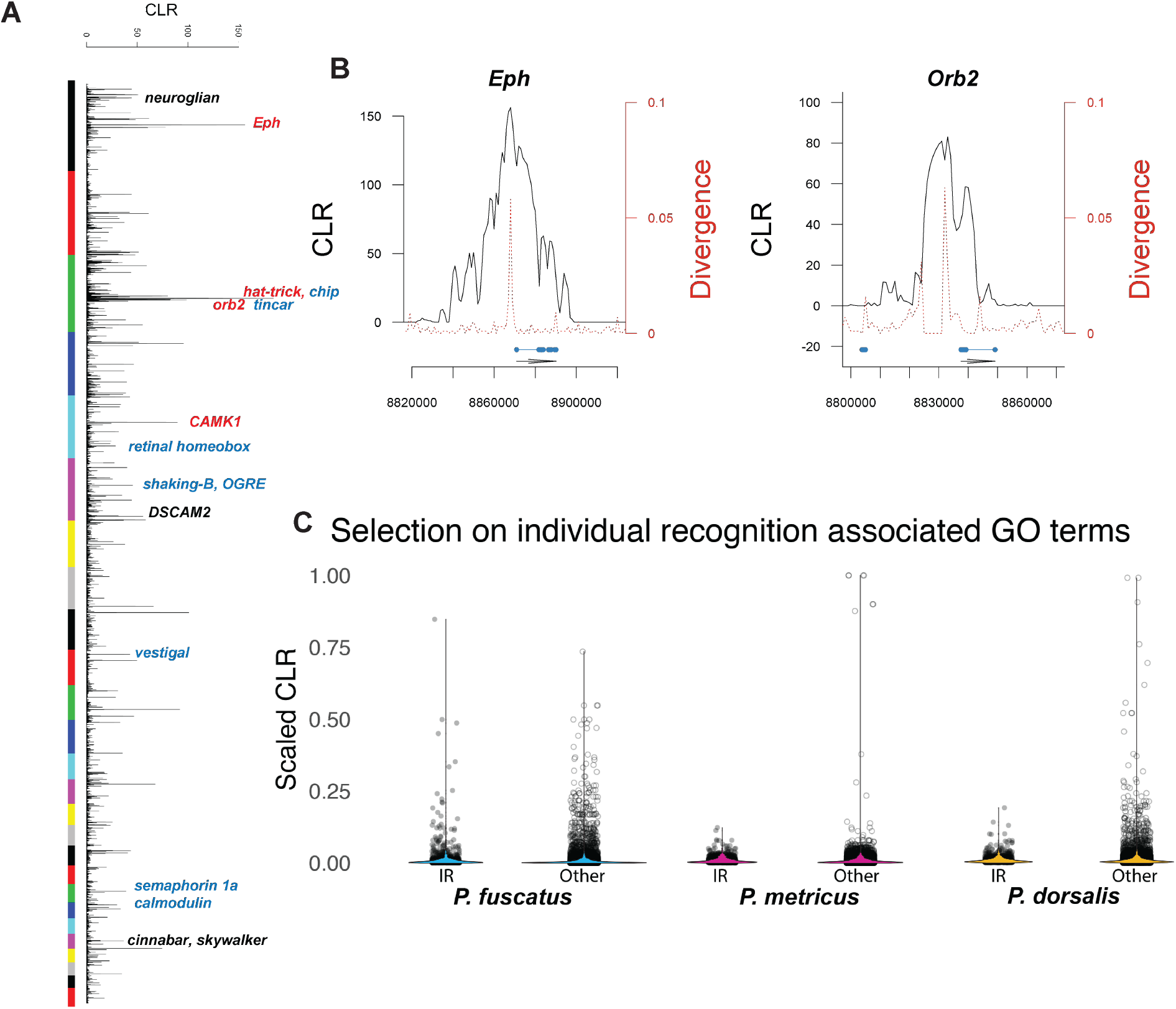
(A) CLR calculated in 1000 bp windows. Colored bars indicate the position of different scaffolds (first 26 shown). Candidate individual recognition genes are shown with annotated function – black: neurogenesis, red: learning and memory, blue: compound eye development. (B) Selected sweeps for two genes involved in the formation of long-term memories, *Eph* and *Orb2*. The sweeps are narrow and patterns of differentiation suggest regulatory elements immediately upstream of the genes are targets of recent selection. (C) Genes in recognition GO categories show evidence of positive selection, though the extent and strength of selection in *P. fuscatus* is far higher. CLR values have been scaled to compare across the three species.

The highest peaks were localized to genes previously connected to cognition or eye development in insects (Figs 2A-B, S6). The most extreme peaks include ephrin receptor tyrosine kinase (*Eph*), which is important for brain development (*18*) and has been experimentally shown to mediate learning and memory in honeybees (*Apis mellifera*) (*19*); Chip (*Chi*) a LIM-homeodomain protein involved in axon guidance (*20*) and compound eye development (*21*); and *Orb2*, which has a role in the development of long-term memory (*22*, *23*). Based on the phenotypes known to have evolved in *P. fuscatus* (*10*, *24*, *25*), genes designated with the Gene Ontology (GO) terms cognition, mushroom body development, visual behavior, learning or memory and eye development were classified as potential candidate genes for individual recognition (hereafter ‘recognition genes’). Overall, approximately 25% of *P. fuscatus* sweeps (36/138) contain candidate recognition genes.

Social selection for recognition abilities may be especially strong in *P. fuscatus*. Alternatively, it may be that genes in recognition-associated GO categories are generally under strong selection in paper wasps and the patterns we observe are unrelated to individual recognition. To distinguish these alternatives, we repeated our analyses looking for recent signatures of selection in *P. metricus* and *P. dorsalis* (Fig. S7-8). We did not observe evidence of recent selection on the homologs of the highly selected genes in *P. fuscatus* in either species. Instead, we find that genes in the top peaks in these species are fatty acid synthases, which create waxes and pheromones, and retrotransposases (Fig S8). To quantitatively assess the difference in selective regimes for recognition genes, we used a mixed effect linear model to compare the relative strength of selection on recognition genes versus all other genes. Across all species, the genes in recognition GO categories had higher CLR values than other genes in the genome (Fig. 2C), suggesting that vision, learning, and memory are often under positive selection in paper wasps. However, there was a strong interaction between gene type and recognition system (Gene Type: df=1, F=6.1, P=0.014; Recognition System: df=1, F=0.03, P=0.9, Gene Type * Recognition System: df=1, F=17.1, P<0.0001). Thus, evidence for recent positive selection on genes involved in cognition and vision is far stronger in *P. fuscatus* than in related species. Similar results were observed when considering specific recognition-associated GO terms. (Fig. S9). However, GO terms predicted to be under selection in all species (e.g. immune response) showed elevated CLR values but did not have a significant interaction with recognition system (Fig. S10). Therefore *P. fuscatus* shows as distinct pattern of strong recent positive selection on genes related to vision, learning and memory. This pattern of selection suggests that social selection for individual recognition has been among the strongest, if not the strongest, selection pressures in *P. fuscatus*’ recent history.

Next, we determined if selection in *P fuscatus* occurred through hard selective sweeps from *de novo* mutations, or soft sweeps from standing variation. We generated a dataset of SNP variation within the Fuscopolistes clade (Fig. S10) by combining re-sequenced data from *P. metricus* and *P. dorsalis* with and additional 17 *P. perplexus* and 18 *P. carolina* re-sequenced genomes, yielding 72 genomes (N = 119 chromosomes). Based on the presence of fixed or nearly fixed *de novo* mutations (Fig S11, Table S4), we classified 46 (33.3%) of the sweeps as hard sweeps and the remainder as soft sweeps. As expected, sweeps were greatly enriched for fixed *de novo* mutations and fixed ancestral polymorphisms (Fig S12, McNemar’s test, p < 0.0001). Multiple candidate recognition sweeps show evidence of classic hard sweeps (e.g. *Eph*, Fig 3A), indicating that evolution of novel cognitive abilities is driven by a mix of hard and soft sweeps (Fig 3A). A similar percentage of sweeps with candidate recognition genes (38.9%; 14/36) were classified as hard sweeps relative to other sweeps (32%, 32/102; z = 0.8, p = 0.4).

**Fig 3.**
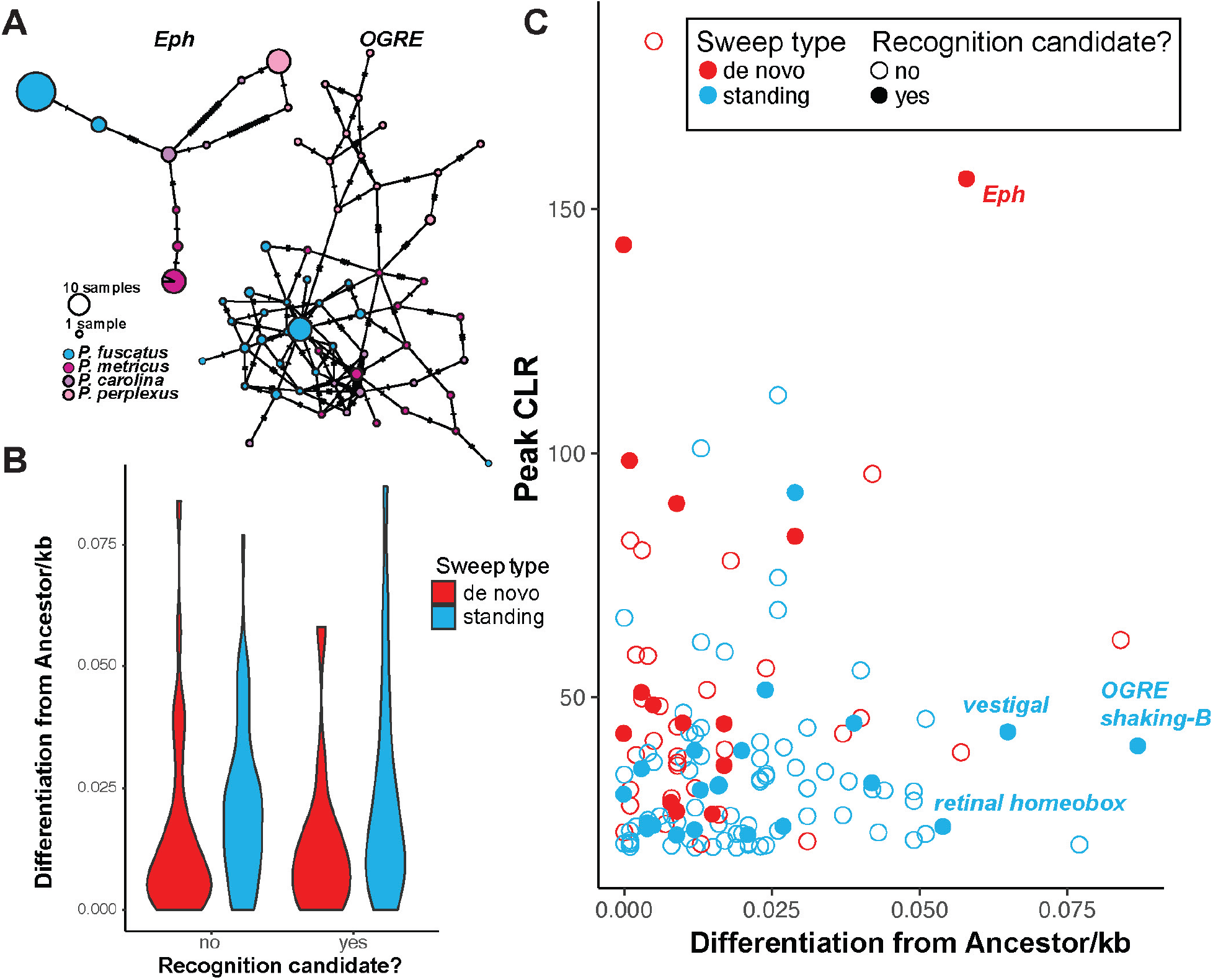
(A) Haplotype networks for *Eph* and *OGRE* associated sweeps. Whereas selection at *Eph* appears to be a hard sweep dominated by one haplotype, the haplotype network at *OGRE* is consistent with a soft sweep with multiple haplotypes. (B) Recognition-associated sweeps with de novo mutations were preceded by selection on standing variation. Violin plots show the Distribution of sweep ages. (C) The relative strength and age for each selected sweep is plotted. Among recognition sweeps, sweeps of standing variation are significantly more prevalent earlier and are involved with eye development. Names of the genes associated with eye development are noted in blue for the oldest sweeps. The oldest hard sweep associated with recognition is associated with the ephrin receptor *Eph*, which mediates long-term memory and mushroom body development.

Evolution of complex novel traits, like social cognition, may be initially biased towards selection on standing variation, or may require novel mutations (*26*). Therefore, we assessed the relationship between candidate recognition sweeps, sweep type, and the relative timing of the selective sweep (Fig 3B, GLM with LRT – Sweep Type: x2 = 74.4, df = 136, p < 0.0001; Recognition candidate: x2 = 0.25, df = 135, p = 0.62; Sweep Type * Recognition: x2 = 7.37, df = 134, p < 0.007). We find that the more divergent (i.e older) sweeps associated with individual recognition are soft sweeps and furthermore are associated with genes for compound eye development. More recently, candidate recognition sweeps are hard selective sweeps and contain genes associated with learning and memory (Fig 3C). These data suggest that the evolution of individual recognition in *P. fuscatus* initially began through improvements in individual discrimination via selection on standing variation in perceptual abilities, followed by strong selection for enhanced cognition. Comparative studies have shown evidence that eye morphology and the relative neural allocation to the optic lobe have recently evolved in *P. fuscatus* (*24*, *25*), consistent with a history of selection on visual abilities.

The extremely strong selection for individual recognition we detect in *P. fuscatus* may be reflective of the role of individual recognition in early season nest-founding associations (*10*), a critical period determining lifetime reproductive success (*27*). A non-mutually exclusive alternative is that novel recognition abilities may generally evolve through intense selection. To assess the generality of our findings, we modeled the co-evolution of individual variability in phenotypes used for recognition and the discrimination and memory abilities of receivers (model details in supplemental materials). We find that when discrimination abilities are initially modest but non-zero, selection gradients for increased discrimination capacity are extreme at the outset of evolving a novel recognition system (Fig 4, S13). As a consequence, recognition systems are predicted to evolve quickly once the benefits of discrimination reach some threshold value. This finding is consistent with our empirical observations. However, nearer the equilibrium value, selection will again be weak, suggesting that observations of weak selection on cognition in wild populations may be the result of these populations being closer to the equilibrium.

**Figure 4.**
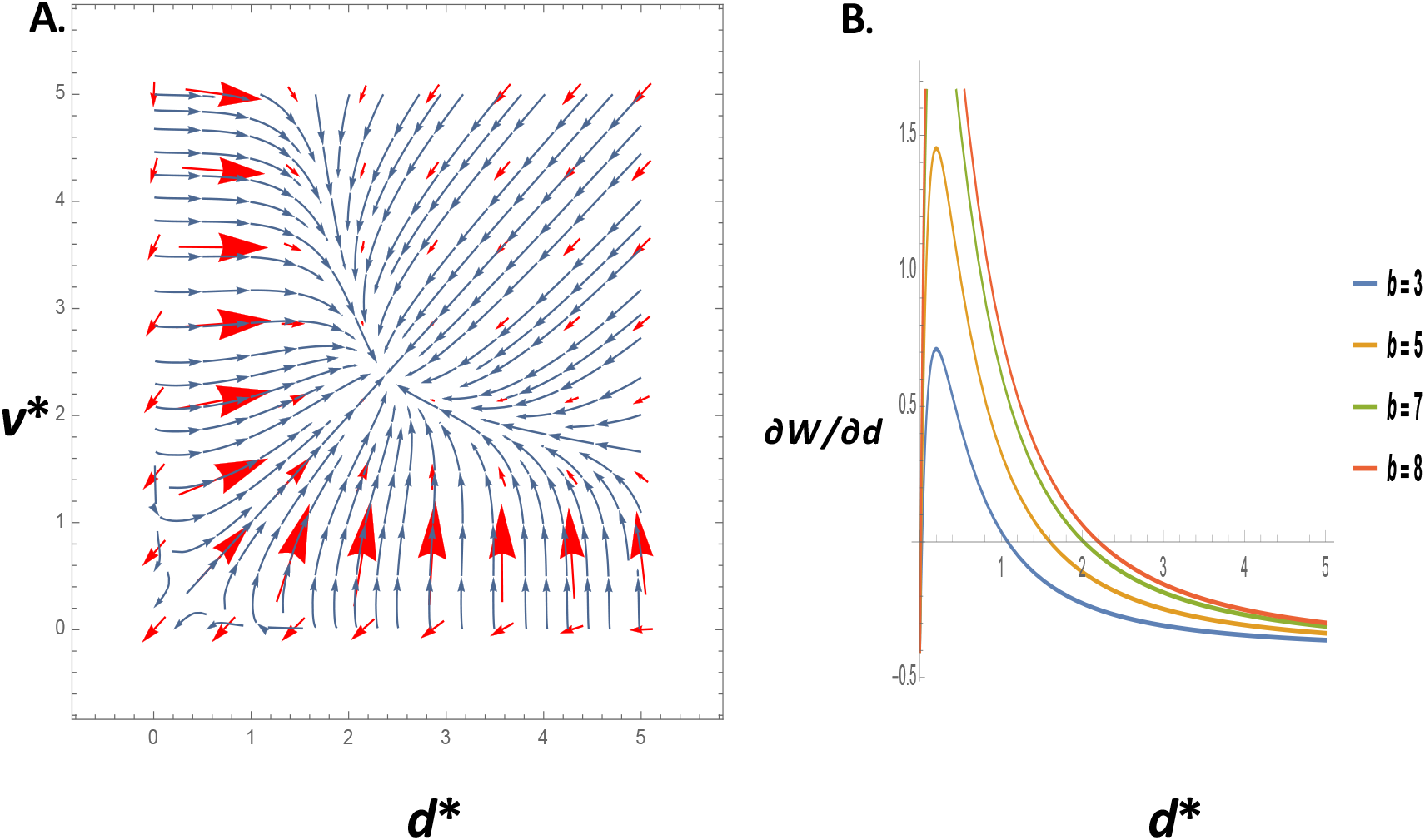
(A) Co-evolutionary dynamics for investment in discrimination ability *d*\* and investment in amplifying cues used in discrimination *v*\*. Red arrows are directional selection gradients; the stream flow-lines are the evolutionary trajectories starting at any given population values of *d*\* and *v*\*. Values of parameters: *a* = 0.80, *b* = 10, *e* = 0.40, and *f* = 0.40. See supplemental materials for model details. (B) Magnitudes of the selection gradient for investment in discrimination as a function of the population investment in discrimination *d*\*, for various values of the benefit *b* of discrimination and the values of *a*, *e*, and *f* assumed in (A), with *v*\* = 2. The strength of selection is highest farther from the equilibrium and for starting values of *d*\* below versus higher than the equilibrium.

The social intelligence hypothesis posits that the cognitive challenges involved in recognizing and tracking individual social relationships selects for improved cognitive abilities (*28*). Recent comparative (*29*) and theoretical (*30*) studies of brain size evolution have called into the question the importance of social interactions as a driver of cognitive evolution. Here we have demonstrated that the evolution of individual recognition, which is the foundation of complex social behaviors, has been associated with intense selection pressures on genes involved in learning, memory and vision. Though social interactions may not be an important factor driving the evolution of intelligence in all taxa, genomic analyses of *P. fuscatus* and our model demonstrate that social interactions can be an extremely potent selection pressure driving cognitive evolution.

## Acknowledgments

This work was supported by an NIH grant (M.S.) and an NSF postdoctoral fellowship (S.E.M). We thank K. Bessler for help with library preparation. T. Blankers, A. Moeller, K. Shaw, and A. Toth provided helpful comments on the project and manuscript. Sequencing data for this project is available at the NCBI Sequence Read Archive under Bioproject accession number PRJNA482994.

